# A comparative study of *in vitro* air-liquid interface culture models of the human airway epithelium evaluating cellular heterogeneity and gene expression at single cell resolution

**DOI:** 10.1101/2023.02.27.530299

**Authors:** Rachel A. Prescott, Alec P. Pankow, Maren de Vries, Keaton Crosse, Roosheel S. Patel, Mark Alu, Cynthia Loomis, Victor Torres, Sergei Koralov, Ellie Ivanova, Meike Dittmann, Brad R. Rosenberg

## Abstract

The airway epithelium is composed of diverse cell types with specialized functions that mediate homeostasis and protect against respiratory pathogens. Human airway epithelial cultures at air-liquid interface (HAE) are a physiologically relevant *in vitro* model of this heterogeneous tissue, enabling numerous studies of airway disease^1–7^. HAE cultures are classically derived from primary epithelial cells, the relatively limited passage capacity of which can limit experimental methods and study designs. BCi-NS1.1, a previously described and widely used basal cell line engineered to express hTERT, exhibits extended passage lifespan while retaining capacity for differentiation to HAE^5^. However, gene expression and innate immune function in HAE derived from BCi-NS1.1 versus primary cells have not been fully characterized. Here, combining single cell RNA-Seq (scRNA-Seq), immunohistochemistry, and functional experimentation, we confirm at high resolution that BCi-NS1.1 and primary HAE cultures are largely similar in morphology, cell type composition, and overall transcriptional patterns. While we observed cell-type specific expression differences of several interferon stimulated genes in BCi-NS1.1 HAE cultures, we did not observe significant differences in susceptibility to infection with influenza A virus and *Staphylococcus aureus*. Taken together, our results further support BCi-NS1.1-derived HAE cultures as a valuable tool for the study of airway infectious disease.

## Introduction

The human airway epithelium is composed of diverse cell types which collectively function to maintain airway integrity, perform mucociliary clearance, and regulate airway immune responses. These include basal cells, the multipotent stem cells which serve as progenitors for other airway epithelial populations^8^, secretory cells (encompassing both goblet cells and club cells), which secrete mucus and other anti-microbial and immunomodulatory peptides^8^, and ciliated cells, which propel directional transport of mucus through coordinated ciliary activity^9^. In addition to these abundant cell types, recent work has identified and highlighted the biological functions of rare cell types which share an airway basal cell precursor, including those of neuroendocrine cells, tuft-like cells, and ionocytes, among others. Neuroendocrine cells are specialized sensory cells that respond to oxygen levels among other diverse chemical and mechanical stimuli and signal these changes to the central nervous system^10,11^. Tuft-like cells are a chemosensory cell population with links to type 2 inflammation through generation of interleukin-25 and leukotriene biosynthesis^12,13^. Ionocytes, present at around 1-2% of airway cellular frequency, have been demonstrated to express high levels of CFTR, the chloride transporter that is dysfunctional in cystic fibrosis, and of other ion transporters^12,14^.

Acknowledging and accounting for this complexity and cellular specialization is important for physiologically relevant study of the airway epithelium in the context of infectious disease. Many respiratory viruses, such as influenza^15^ and SARS-CoV-2^16^ infect ciliated cells. Human strains of Influenza A virus (IAV) utilize α2,6-linked sialic acid (SA) as their receptor which is expressed in a graded fashion along the upper airway, predominantly in ciliated cells and to a lesser extent, goblet (secretory) cells^17,18^. In contrast, avian influenza strains (H5N1) utilize α2,3-linked SA which is expressed largely in alveolar type II cells in the lower respiratory tract^18^. For SARS-CoV-2, ciliated cells express both the viral receptor ACE2 and protease TMPRSS2 required for fusion at the cell membrane, and motile cilia have been shown to be required for efficient replication in *in vitro* airway epithelial models^9,16^. Airway mucus, produced by secretory cells, is also critical for innate defense against respiratory infections. *Staphylococcus aureus* (*S. aureus*), *Pseudomonas aeruginosa* (*P. aeruginosa*) and *Haemophilus influenza* (*H. influenza*) have all been shown to upregulate the production of mucins, a process critical for trapping invading pathogens which can also become detrimental in patients with respiratory disease^19–21^. Experimental model systems which account for these factors are important for enabling a comprehensive understanding of disease caused by respiratory pathogens.

Primary differentiated polarized human airway epithelium (HAE) cultures at air-liquid interface are an established, physiologically relevant *in vitro* system used to study mechanisms of airway homeostasis and disease^1,6,7,22^. HAE cultures are classically derived from normal human bronchial epithelium (NHBE) acquired via airway biopsy, which are differentiated into pseudostratified epithelia when grown on Transwells at an air-liquid interface^1^. HAE cultures recapitulate many features of the human airway epithelium, including characteristic airway cell types and secreted extracellular environment^2,3^, and they have been widely used in airway research ^1,3^. A limitation of HAE cultures derived from primary precursor cells is their restricted passaging capacity before losing differentiation potential or becoming senescent^2^. This complicates the design of larger experiments, as sufficient material may require multiple donor sources, thereby introducing confounding genetic variation and increasing cost. Although achieved in some studies^6,23^, limited passage capacity also poses feasibility challenges to genetic manipulation of HAE cultures, which can require extended, multi-passage selection of primary cell populations.

Despite the technical barriers to genetic engineering of HAE cultures, primary bronchial basal epithelial cells can be transgenically engineered to express hTERT, prolonging the proliferative capacity of HAE culture precursor cells. BCi.NS1.1 is an airway basal cell line isolated from a healthy, non-smoking male donor, engineered to express hTERT, which extends passage capacity to up to 40 times without losing differentiation potential, as compared to a limit of only 3-4 passages with untransduced primary cells^5^. Previous studies have demonstrated that BCi-NS1.1 differentiates into HAE cultures with similar morphology and cell type composition to primary HAE cultures, and this system has been leveraged for a variety of applications^5,24–26^. However, the effects of hTERT expression on rare airway cell composition and associated gene expression programs in differentiated HAE cultures have been incompletely characterized, and any potential functional consequences for infection models with respiratory pathogens have not been directly explored. Furthermore, while hTERT-derived HAE cultures should exhibit less culture-to-culture variation than HAE cultures derived from different donors, this has not been directly assessed.

Here, we perform an in-depth comparison of HAE cultures derived from BCi-NS1.1 cells (n = 3 independent differentiations) and HAE cultures derived from independent, commercially available donor primary cells (n = 3). Initially, we evaluated culture microarchitecture and canonical human airway epithelial cell type markers by histology and immunohistochemistry. We then used single cell RNA-Sequencing (scRNA-Seq) for comprehensive and unbiased assessment of constituent cell populations, differentiation states, and gene expression programs. To assess potential functional differences relevant to airway infectious disease, we evaluated HAE culture responses to infection with two common respiratory pathogens, influenza A virus (IAV) and *Staphylococcus aureus* (*S. aureus*). Overall, we sought to thoroughly define the similarities and differences of hTERT-expressing BCi-NS1.1-derived HAE cultures and primary cell-derived HAE cultures to inform appropriate applications of these powerful *in vitro* experimental models.

## Methods

### HAE cultures

BCi-NS1.1 cells (passage 17-25) (kind gift of Dr. Ronald Crystal) and Normal Human Bronchial Epithelial cells (NHBE) (cat. no. CC-2541, Lonza, lot numbers 623950, 626776, and 630564) were seeded in Pneumacult™-Ex Plus Medium (StemCell) and passaged at least two times before plating (7×10^4^ cells/well) on rat-tail collagen type-I coated permeable Transwell membrane supports (6.5 mm diameter, 0.4 μm pore size; Corning Inc) to generate HAE cultures. HAE cultures were maintained in Pneumacult™-Ex Plus Medium until confluent, then grown at air-liquid interface with Pneumacult™-ALI Medium in the basal chamber for approximately 3 weeks to form well-differentiated, polarized cultures.

### Histology, immunohistochemical staining and imaging

Embedding, sectioning and Hematoxylin and Eosin (H&E)/Periodic acid-Schiff (PAS) Alcian blue staining of HAE cultures were performed as described previously by our laboratory^27^. For immunohistochemical staining and imaging of HAE cultures, prepared slides were deparaffinized in Xylene (cat. no. 301-038-4000, Crystalgen) for 30 minutes, then rehydrated in a graded series of ethanol dilutions in water. Antigen unmasking was performed by heating slides submerged in 1X Citrate buffer (pH 6.0) (cat. no. ab64214, Abcam) in a microwave for 1 minute at 100% power, then for 8 minutes at 10% power. Slides were cooled to room temperature, washed twice with BupH Tris Buffered Saline (cat. no. 28376, Thermo Fisher) 0.05% Tween-20 (cat. no. BP337-500, Thermo Fisher), and blocked with Pierce™ SuperBlock™ T20 (TBS) Blocking Buffer (cat. no. 37581, Thermo Fisher) 0.5% Triton X-100 (cat. no. 85111, Thermo Fisher) for 30 minutes. Blocking buffer was discarded and slides were incubated with primary antibody diluted in blocking buffer overnight at 4°C. Primary antibodies included α-villin-1 diluted 1:500 (cat. no. NBP1-85335-25ul, Novus Bio), α-cytokeratin 5 diluted 1:500 (cat. no. Ab75869, abcam), α-CC10 diluted 1:400 (cat. No. 365992, Santa Cruz), α-MUC5B diluted 1:2000 (cat. no. PA5-82342, Thermo Fisher), and α-MUC5AC diluted 1:1000 (cat. no MA5-12178, Thermo Fisher). Slides were washed three times with wash buffer, then incubated in secondary antibody diluted in blocking buffer for one hour at room temperature. Slides were washed twice with wash buffer and three times with water, then mounted with coverslips using mounting medium from Pierce™ Peroxidase IHC Detection Kit (cat. no. 36000, Thermo Fisher). Slides were imaged using a Keyence BZ-X810 Fluorescent Microscope (cat. no. BZ-X810, Keyence) and images were analyzed using BZ-X800 Analyzer.

### Cell type quantification by immunofluorescence

Immunofluorescence images of the villin-1, KRT5, CC10, MUC5AC and MUC5B labeled HAE cultures were quantified using Imaris microscopy image analysis software version 9.8.2. The stitched image depicting the entire length of each HAE culture transection for each cell type stain was used for image quantification. Two technical replicate images of each biological replicate BCi-NS1.1 or primary HAE culture were used for image quantification. For each marker, signal area was determined using the surfaces creation tool and setting the intensity threshold to exclude any background as determined by HAE culture controls labeled without primary antibody. The total area of phalloidin staining was used as the total area of each HAE culture transection, and the percentage occupancy of cell type marker is represented as a percentage of this phalloidin stained area.

### Isolation and preparation of single cell suspensions from HAE cultures for scRNA-Seq

HAE cultures maintained for 3 weeks post-airlift were processed for scRNA-Seq. Single cell suspensions of HAE cultures were prepared as described previously^27^. To enable multiplexing and doublet detection, cells were labeled with barcoded antibodies described previously^28^. Briefly, approximately 200,000 cells per sample were resuspended in staining buffer (PBS, 2% BSA, and 0.01% Tween) and incubated for 10 minutes with Fc block (TruStain FcX (BioLegend) and FcR blocking reagent (Miltenyi). Cells were then incubated with oligonucleotide-conjugated hashing antibodies (generated in-house by the New York Genome Center as described^29^) for 30 min at 4°C. After labelling, cells were washed 3 times in staining buffer. After the final wash, cells were resuspended in PBS plus 0.04% BSA, filtered, and counted. Cells were pooled (6 samples per pool), super-loaded to the 10X Genomics Chromium Controller (~4,170 cells per sample, ~25,000 cells per lane) and processed with the Chromium Next GEM single-cell 5′ library and gel bead kit according to manufacturer’s protocols. Hashtag-oligonucleotide (HTO) additive oligonucleotide primer was spiked into the cDNA amplification PCR, and the HTO library was prepared as described previously^29^.

### scRNA-Seq analysis

#### Read mapping and quantification

CellRanger count (version 6.1.2, 10X Genomics Inc.) was used to assign cell barcodes and map reads to transcriptome reference (GRCh38-2020-A transcriptome reference) with default parameters. HTO quantification was also performed using CellRanger count via feature barcode detection.

#### Data preprocessing and integration

scRNA-Seq analyses were performed in R (v4.0.3) using *Seurat* 4.0^30^. Samples were demultiplexed by hashtag oligo (HTO) counts produced from CellRanger count using the *HTODemux* command in Seurat with default parameters. Demultiplexing quality control (QC) was assessed by inspection of a dimensionality reduction plot of HTO counts by t-SNE. Cell barcode × feature (gene) count matrices were filtered to exclude cells with >18.7% mitochondrial gene content (dead or dying cells) or those with feature (gene) counts outside of the interval from 1,505-5,857 or unique molecular identifier (UMI) counts outside of the interval from 4,569-24,992, which correspond to 1.2 and 1.5 standard deviations from the mean number of counts (log), respectively. Intra-hash heterotypic doublet detection and removal were performed with scDoubletFinder (v1.4.0)^31^. Gene expression data were normalized with scTransform (v.0.3.3)^32^ using top 3,000 variable genes and including a regression factor for “cell cycle difference,” determined by subtracting the G2 / M score from the S score for each cell calculated using the gene sets described by Tirosh and colleagues^33^. To minimize the effect of donor-specific variation on cluster assignment, data integration was performed across all replicates using 2,000 gene features and Seurat commands *PrepSCTIntegration*, *FindIntegrationAnchors* and *IntegrateData*.

#### Data clustering and annotation

Principal component analysis (PCA) was performed on normalized, integrated data, and the first 30 principal components (PCs) were used for clustering and nearest neighbor detection. An iterative approach was used to remove low quality clusters of cells: initial clustering was performed using the graph-based Louvain algorithm with multilevel refinement^34^ and a resolution parameter of 1.5. Two clusters (333 cells total) were excluded from downstream analysis due to low UMI counts and high mitochondrial content, respectively. Data were then re-clustered and assessed for stability at a range of resolutions using *Clustree* (v.0.5.0)^35^. A final resolution parameter of 1.4 was selected from these results. Clusters were annotated to canonical airway cell populations based on expression of established marker genes^3,11,12,14^. Cell annotations were verified by comparing assignments to predictions obtained by label transfer from a recently published scRNA-seq dataset of similar HAE cultures^36^.

#### Pseudobulk differential gene expression analyses

Differential gene expression analyses were conducted on “pseudobulk” gene expression profiles using *edgeR* (v3.32.1)^37^ and *scran* (v1.18.7)^38^, a strategy found to provide better type I error control than individual cell-based differential expression workflows^39^. Briefly, per cell gene counts were summed to aggregate “pseudobulk” profiles for each cell population replicate, with a minimum threshold for inclusion of 25 cells. Gene expression linear models were constructed for each cell population with at least two samples per condition (BCi-NS1.1, primary) passing inclusion thresholds. Differential gene expression was defined as an adjusted p-value < 0.05 (Benjamini-Hochberg) and an estimated log2 fold-change > 1. Additionally, genes expressed in fewer than 10% of cells in any pseudobulk sample were excluded from differential expression tests.

#### Pseudobulk gene set enrichment analyses

Gene set enrichment analyses were performed using CAMERA^40^ for the molecular signatures database (MSigDB)^41^ C2 canonical pathways gene set (n = 3,050) and HALLMARK gene set (n = 50) collections, with the edgeR linear models described in section 2.6.4. Significant set enrichment was defined as an adjusted p-value < 0.05.

#### Pseudotime trajectory inference

Cell differentiation trajectories were inferred using *Slingshot*^42^. Analyses included only cells along the basal-secretory axis. Principal curves were fit in integrated UMAP space and rooted to the basal cell cluster. Differential progression testing was assessed using a Kolmogorov-Smirnov Goodness of Fit Test between conditions using the *progressionTest* command with default arguments.

#### Differential abundance testing

Cell abundance was modeled using *edgeR* as described^43^, without normalization for total cell count. Significant differential abundance was defined as any change in cell population frequency at a p-value < 0.05.

#### Infection of HAE cultures with influenza A virus

HAE cultures were washed apically twice with room temperature PBS, then 5×10^6^ PFU of influenza A/California/07/2009 virus (BEI resources, NR561 13663) was added apically in 50 μl of PBS and incubated for one hour at 37°C. PBS was aspirated after 1 hour of infection and HAE cultures were washed apically once with room-temperature PBS, then incubated at 37°C for 24 hours. Mock-infected cultures were treated with the same protocol as IAV-infected cultures but using only PBS. After 24 hours, cultures were harvested for either quantification of viral PFU, RT-qPCR, LDH quantification, imaging, or western blot analysis. For quantification of viral PFU, a 200μl apical wash was collected from the cultures and frozen at −80°C for plaque assay on MDCK cells. For RT-qPCR, cultures were washed once, and the membrane was cut out and submerged in buffer RLT then frozen at −80°C for RNA extraction using RNeasy Kit (cat no. 74104, Qiagen), cDNA synthesis using SuperScript™ III First-Strand Synthesis System (cat no. 18080051, Thermo Fisher) and RT-qPCR using PowerUp™ SYBR™ Green Master Mix (cat no. A25741) and RPS11 as a housekeeping gene. For LDH quantification a 200μl apical wash was collected from the cultures and LDH release was measured in 50μl of apical wash in triplicate using CyQUANT™ LDH Cytotoxicity Assay (cat. no. C20300, Thermo Fisher). For western blot analysis, cultures were washed once and the cells were lysed by adding 200μl of 1x Invitrogen™ 4X Bolt™ LDS Sample Buffer (cat. no. B0007, Fisher Scientific). Lysate was frozen at −20°C for western blot analysis. For imaging, cultures were submerged in 4% PFA, incubated overnight at 4°C, then transferred to PBS. Staining and imaging of infected cultures was performed as described previously by our laboratory^27^.

#### Infection of HAE cultures with S. aureus

*Staphylococcus aureus* (*S. aureus*) USA300 strain AH-LAC^44^ was routinely grown at 37°C on tryptic soy agar (TSA). For infection of HAE cultures, overnight bacterial cultures were grown in 5 mL tryptic soy broth (TSB) in 15-mL conical tubes under shaking conditions at 180 rpm with a 45° angle. A 1:100 dilution of overnight culture was subcultured into TSB and incubated for another 3 hours. Bacteria was washed once with 5 mL PBS then diluted using Optical Density 600 (OD600) measurements to an OD of 0.8, corresponding to a concentration of 2×10^7^ CFU/mL in PBS. HAE cultures were washed apically twice with room temperature PBS, then 1×10^6^ CFU of AH-LAC was added apically in 50 μl of PBS and incubated for one hour at 37°C. PBS was aspirated after 1 hour of infection and HAE cultures were washed apically three times with room-temperature PBS, then incubated at 37°C for 18 hours. After 18 hours, cultures were harvested for either quantification of bacterial CFU, LDH quantification or imaging. For quantification of bacterial CFU, the membrane was cut out and submerged in 1% Saponin to lyse cells, and lysate was serially diluted and plated on TSA. Samples were harvested and prepared for LDH quantification and imaging as described above.

#### Treatment of HAE cultures with IFNβ

For treatment with IFNβ, basal media was removed from the cultures and media containing 500 U/mL human IFNβ (cat. no. IF014, lot 3481402, Millipore Sigma) was added to the basal chamber. IFN-treated cultures were incubated for 24 hours at 37°C. After 24 hours, IFNβ-treated cultures were harvested and prepared for either RT-qPCR or western blot analysis as described above.

## Results and Discussion

To assess similarities and differences between HAE cultures differentiated from the hTERT-expressing bronchial basal epithelial cell line BCi-NS1.1^5^ and unmodified, primary-derived precursor cells, we generated HAE cultures from three donors of primary normal human bronchial epithelial cells (NHBE), as well as three biological replicates of HAE cultures from BCi-NS1. 1^5^ (Fig. 1a).

**Figure 1:**
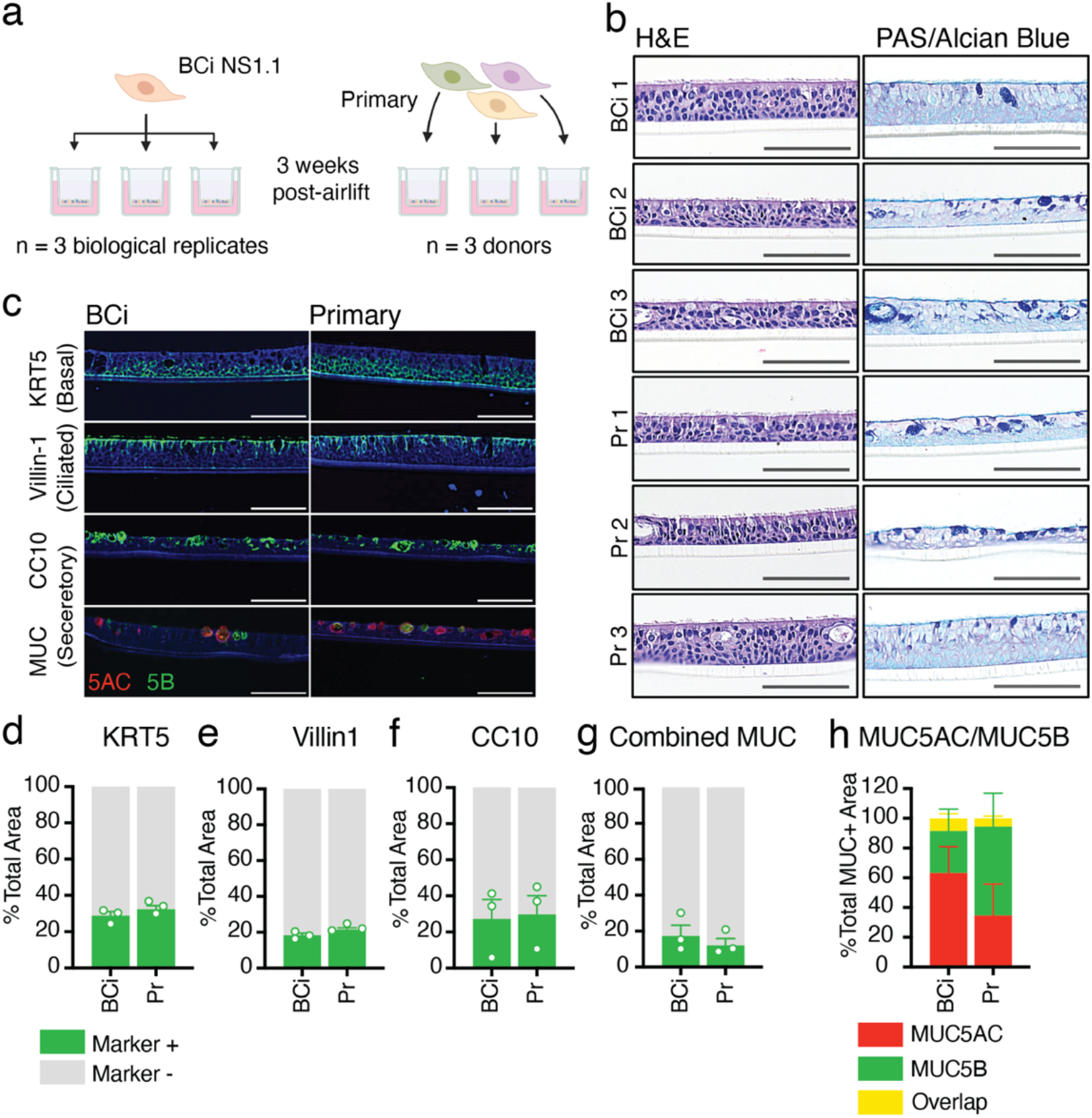
Analysis of epithelial structure and cell types produced by BCi-NS1.1- and primary-derived HAE cultures. **a**. Diagrammatical representation of experimental design. HAE cultures were generated from either primary cells or the hTERT-expressing precursor cell line BCi-NS1.1. Cells were differentiated on Transwells for three weeks at air-liquid interface to generate three replicates of BCi-NS1.1-HAE cultures (derived from one donor), and three HAE cultures from three separate primary donors. **b**. Hematoxylin & Eosin (H&E) and Periodic Acid-Schiff (PAS)-Alcian blue staining of formalin-fixed paraffin-embedded transections of HAE cultures from either BCi-NS1.1 or primary precursor cells. All cross-sectional images are oriented with the basolateral surface of the culture on the bottom of the image and the apical surface on the top of the image. Scale bars = 150μm **c**. Cell type-specific immunohistochemical staining of transections of BCi-NS1.1- or primary-derived HAE cultures. DAPI (nuclei) stained in blue. Cytokeratin 5 (KRT5, basal cells), villin-1 (ciliated cells), CC10 (encoded by SCGB1A1) (secretory cells) and MUC5B (secretory cells) stained in green. MUC5AC (secretory cells) stained in red. Scale bars = 150μm. **d-g**. Quantification using Imaris software of cells stained positive for each cell-type specific marker, KRT5 (c), villin-1 (d), CC10 (e), and MUC (f) (sum of MUC5B and MUC5AC positive cells), represented as a percentage of the total phalloidin (actin)-stained area of the HAE cultures. **h**. Quantification using Imaris software of cells stained positive for MUC5B, MUC5AC, or both, represented as a percentage of total MUC+ cells.

### Both primary and BCi-NS1.1 precursor cells differentiate into pseudostratified epithelia that include basal, secretory, and ciliated cells

We first characterized the histological morphology of HAE cultures derived from BCi-NS1.1 cells and primary cells. Hematoxylin and eosin (H&E) staining of HAE culture sections demonstrated that cultures derived from both BCi-NS1.1 and primary precursors adopt a polarized pseudostratified epithelia morphology with ciliated cells facing the apical side (Fig. 1b). PAS-Alcian blue staining revealed the presence of intracellular mucins (dark blue), suggesting the presence of secretory cells (Fig 1b). Overall, individual cultures exhibited subtle differences in epithelium thickness, secretory cell frequency, and secretory cell hyperplasia, but these differences were not associated with precursor cell source.

Next, we determined the presence and intra-epithelial organization of ciliated, basal, and secretory cells by immunofluorescence microscopy. Cultures were immunolabeled for canonical cell type-specific markers: villin-1 for ciliated cells^45^, cytokeratin-5 (KRT5) for basal cells^4^, and CC10 (encoded by the *SCGB1A1* gene), MUC5B and MUC5AC for secretory cell^1–7^.s^3^. We observed KRT5 expression in small, brick-shaped cells lining the basolateral surface consistent with basal cells (Fig. 1c, Fig. S1a). Image quantification of this label identified KRT5-positive cells to account for approximately 31% of total cell area in both BCi-NS1.1- and primary-derived HAE cultures (Fig. 1d). We observed villin-1 in columnar cells lining the apical surface, consistent with ciliated cells (Fig. 1c, Fig. S1b). These cells accounted for approximately 20% of total cell area in both BCi-NS1.1- and primary-derived HAE cultures (Fig. 1e). Lastly, secretory cell makers were present in heterogeneous cells, including large globular cells facing the apical surface (Fig. 1c, Fig. S1c). CC10-positive cells made up on average 28% of total cell area in both BCi-NS1.1- and primary-derived HAE cultures, although the number of cells positive for this marker was more variable between cultures than either KRT5 or villin-1 (Fig. 1f). Cells positive for either MUC5AC or MUC5B generally made up less than 20% of the total cell area (Fig. 1c,g, Fig. S1d). While technical constraints prohibited multiplex staining for CC10 and mucins, dual staining for MUC5B and MUC5AC revealed that most but not all MUC-positive cells expressed one of these two mucins but not both (Fig. 1c,g, Fig. S1d). Although variable across cultures, the proportions of MUC5B vs MUC5AC positive cells (Fig. 1g, Fig. S1d) were not associated with precursor cell source. Taken together, immunofluorescence imaging of cell-type-specific markers revealed that ciliated, basal, and secretory cells are present in both BCi-NS1.1 and primary cultures at similar frequencies.

### scRNA-Seq resolves constituent cell populations of in vivo airway epithelium

To assess the cellular composition at high resolution along with associated gene expression patterns of primary and BCi-NS1.1-derived HAE cultures, we performed scRNA-Seq. In total, we analyzed 12,801 single cells (post quality-control filtering) from three samples of each HAE culture source (n = 3 independent differentiations of BCi-NS1.1, n = 3 different donors of primary cells). Unsupervised graph-based clustering identified 12 airway epithelial cell populations,, which were annotated based on RNA expression of canonical marker genes (Fig. 2A) and label transfer predictions from a recently reported scRNA-Seq analysis of HAE^36^. scRNA-Seq data included basal cells (*KRT5*^+^, *TP63*^hi^), proliferating cells (*MKI67*^+^, *TOP2A*^+^), suprabasal cells (KRT6A^+^, KRT16^+^), “intermediate” (i.e. along the basal-to-secretory differentiation axis) cells, three populations of secretory cells (I: *MUC5AC*^lo^ *MUC5B*^lo^, II: *MUC5AC*^lo^ *MUC5B*^hi^, and III: *MUC5AC*^hi^ *MUC5B*^lo^), deuterosomal cells (*DEUP1*^+^, *FOXN4*^+^), ciliated cells (*FOXJ1*^+^, *CAPS*^+^), tuft-like cells (*LRMP*^+^, *ASCL2*^+^), ionocytes^12,14^ (*FOXI1*^+^, *CFTR*^+^), and neuroendocrine cells (*ASCL1*^+^, *HOXB5*^+^)(Fig. 2b). Cell annotation was consistent across samples (Fig. S2a). The relative frequencies of most cell populations were similar between primary and BCi-NS1.1-derived HAE cultures, with the exception of MUC5AC^hi^ secretory III cells, deuterosomal cells, and ciliated cells, which were more highly represented in BCi-NS1.1-derived HAE cultures (Fig. 2c). Notably, both culture sources exhibited rare ionocytes, tuft-like cells, and neuroendocrine cells at similar frequency. Unexpectedly, we detected far fewer ciliated cells in scRNA-Seq data than in IF (~2% vs ~20%). Based on our prior experience analyzing HAE by scRNA-Seq, we attribute this discrepancy to technical issues during sample processing resulting in the selective loss of ciliated cells prior to 10X Chromium loading. With severely reduced ciliated cell numbers, we focused the remainder of our scRNA-Seq analyses on robustly sampled basal and secretory cells. Despite these limitations, scRNA-Seq identified the expected constituent cell populations of the airway epithelium with few differences in frequency across culture types.

**Figure 2:**
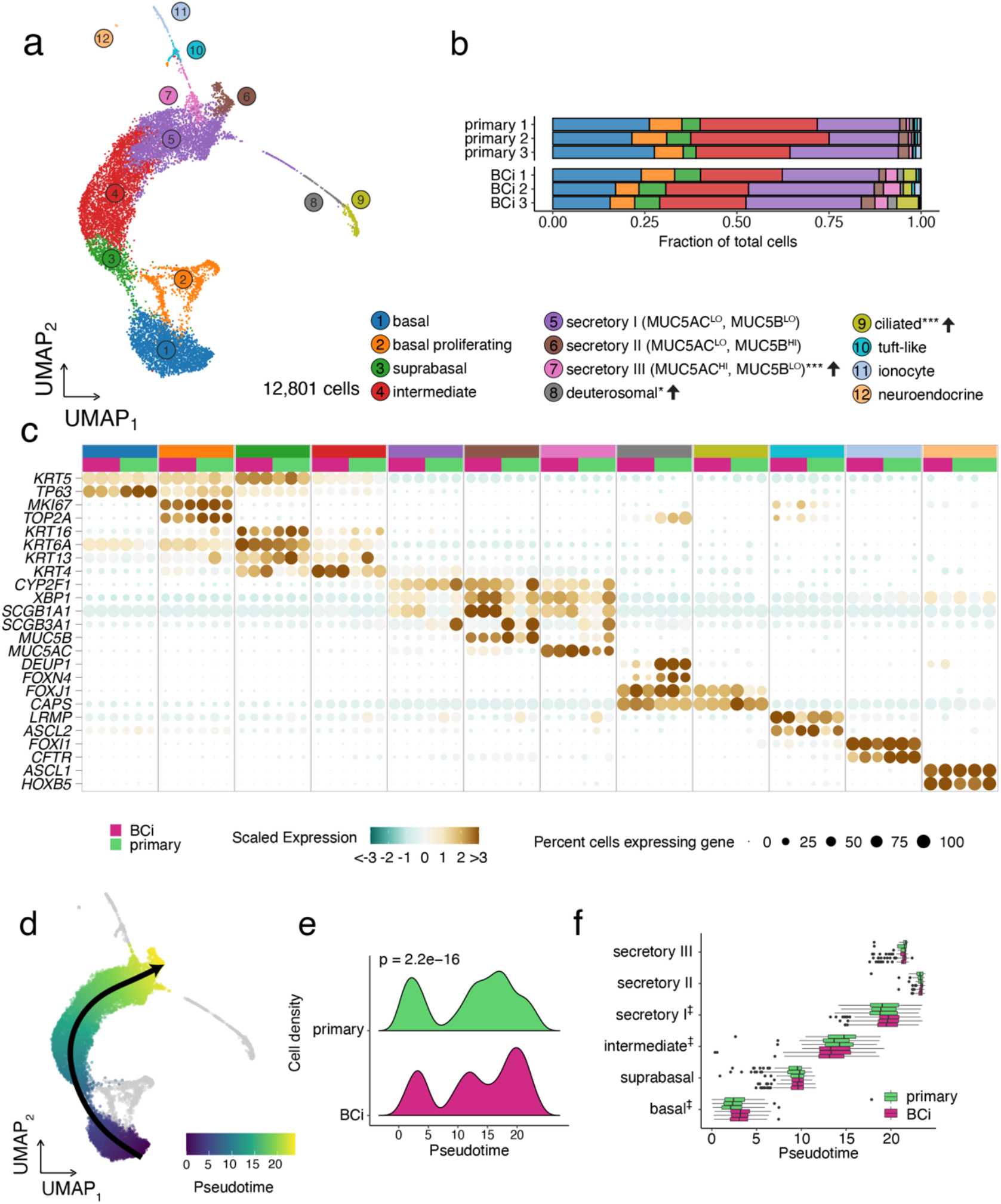
Cell population annotation and pseudotime analyses of scRNA-Seq data from BCi-NS1.1 and primary –derived HAE cultures. **a**. A UMAP dimensionality reduction plot of 12,801 single cell transcriptomes, colored by cell population assignment. Semi-supervised annotation of scRNA-Seq data resolved 12 HAE cell populations: basal cells, proliferating cells, suprabasal cells, intermediate cells, three populations of secretory cells differentiated by their MUC5AC/B expression, ciliated cells, and rare deuterosomal cells, tuft-like cells, ionocytes, and neuroendocrine cells. **B**. Cell proportions by biological replicate. Cell populations with altered frequencies are noted in the legend indicating the change in BCi-NS1.1 relative to primary HAE cultures along with the significance level (* P < 0.05, ** P < 0.01, *** P < 0.001). **c**. A dot plot summarizing the scaled expression (dot color) and level of expression (dot size) of signature marker genes for each cell population, broken out by biological replicate. **d**. Pseudotime trajectory inference identified a single lineage spanning basal-to-secretory cells (both HAE culture types superimposed). Proliferating cells and low frequency cell types (grey) were excluded from trajectory analysis. **e**. Cell densities along the basal-to-secretory trajectory reveal differential cell progression between the BCi-NS1.1- and primary cultures (Kolmogorov-Smirnov Goodness of Fit Test, p = 2.2e-16). **f**. A boxplot of cell pseudotime values by cell type and scRNA-Seq replicate. Hypothesis testing for differential median pseudotimes for basal, intermediate and secretory I cells all reached the minimal p value of 0.1 via a Wilcoxon Rank Sum Test with 3 replicates per group (‡) but do not clear a 95% significance level.

### Pseudotime analysis of scRNA-Seq data reveals a common basal-to-secretory cell differentiation trajectory but differences in distribution of BCi-NS1.1-derived HAE cultures

As we observed subtle differences in the frequency of cell populations between primary- and BCi-NS1.1-derived HAE cultures, we investigated the possibility that these differences in cellular composition could be related to the airway cell differentiation process. To this end, we inferred differentiation trajectories by pseudotime analysis. We restricted our analysis to basal, suprabasal, intermediate, and secretory populations based on the established basal-to-secretory differentiation path^3^ and to eliminate potential confounding factors from actively proliferating cells and rare cell types for which we lacked robust sampling. A single lineage was identified for both culture types: basal-to-suprabasal-to-intermediate-to-secretory (I, III, II) (Fig. 2d). Differential distribution testing indicated significant differences in progression along this trajectory between the culture conditions, with primary-derived HAE cultures exhibiting relatively more cells in basal and “early” secretory states in comparison to the BCi-NS1.1-derived cultures (Fig. 2e). A closer inspection of the pseudotime distribution for each cell population illustrated several shifts in the median pseudotime values: in comparison to primary-derived HAE cultures, the median basal and secretory I pseudotime values were higher in BCi-NS1.1-derived HAE cultures, while the median pseudotime value for intermediate cells was lower (Fig. 2f). This suggests that, while general cell population abundances are consistent across culture types, there exist modest but significant differences in the distribution of BCi-NS1.1-derived HAE cultures along the basal-to-secretory differentiation axis.

### HAE cultures exhibit generally similar gene expression patterns by cell population across BCi-NS1.1- and primary-derived cultures

We next assessed gene expression patterns in each cell population to compare HAE cultures from the two culture progenitors. “Pseudobulk” gene expression profiles (by cell population and replicate) were aggregated and clustered by Spearman correlation distance (Fig. 3a). Data clustered primarily by major cell population (i.e. basal, secretory, ionocyte, ciliated/deuterosomal), and pairwise comparisons further emphasize similarities across the different culture progenitors by cell population. Interestingly, BCi-NS1.1-derived ‘intermediate’ cell populations clustered with BCi-NS1.1-derived suprabasal cell populations, a result consistent with pseudotime analysis. Similarly, principal component analysis (PCA) revealed a strong segregation by cell population in the first three PCs (accounting for 72.6% total variation), with basal/ciliated, basal/secretory, and ionocyte divergence captured by the first three components, respectively (Fig. 3b). PC4 appeared to segregate data by culture type but described a relatively small amount (6.1%) of total variation. Taken together, these results indicate that by gene expression, cell populations are largely similar across BCi-NS1.1 and primary HAE cultures, with a relatively small amount of variation associated with differences in the culture progenitor groups.

**Figure 3:**
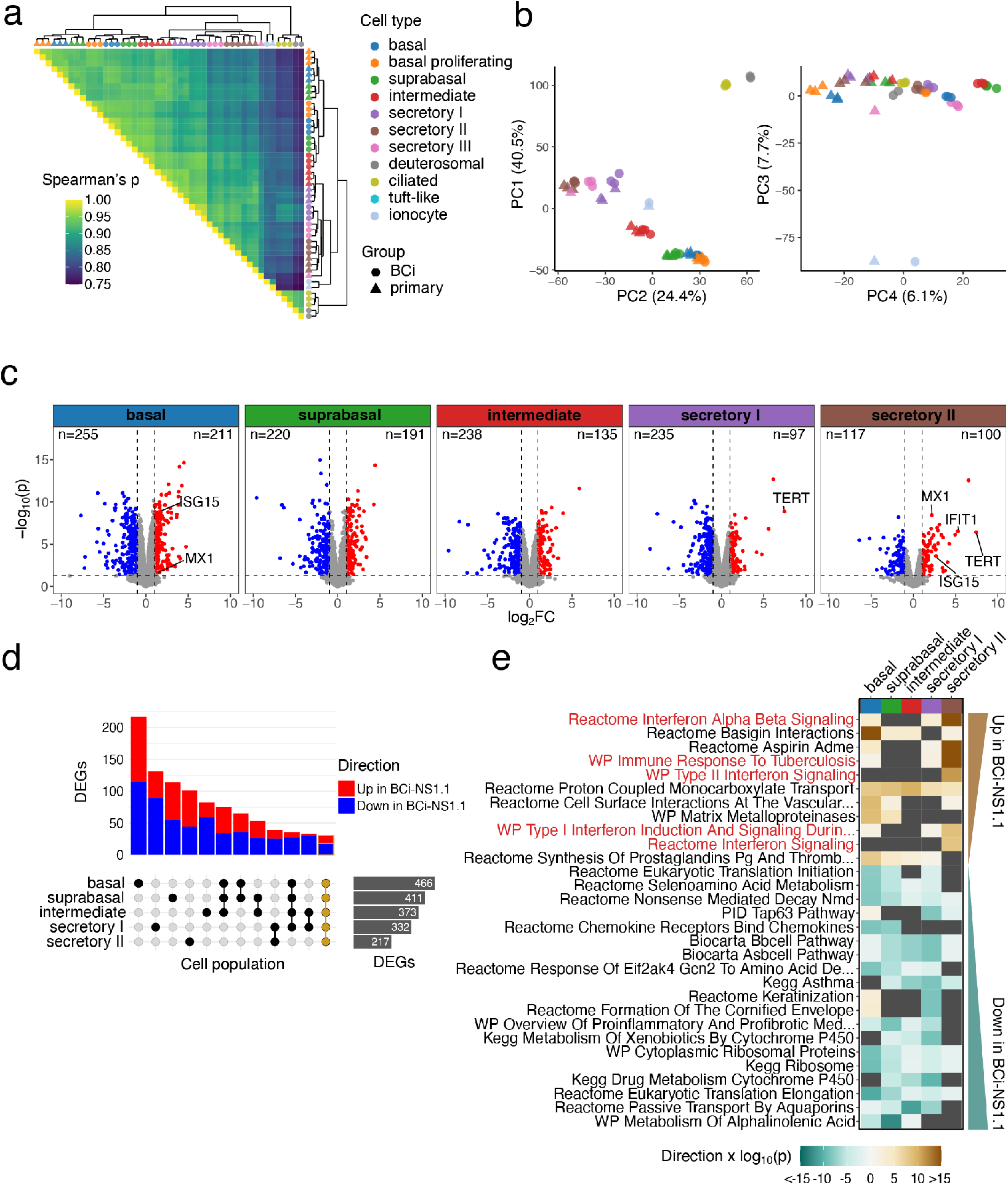
Gene expression analyses of BCi-NS1.1 and primary –derived HAE cultures. **a**. Triangular heatmap of Spearman correlation coefficients for pseudobulk transcriptional profiles aggregated by cell population (color) and culture group (shape) scRNA-Seq replicate. Row and column clustering was determined by Ward’s linkage method and shown as a marginal dendrogram. **b**. Principal component analysis (PCA) of pseudobulk transcriptional profiles, showing point plots for principal component (PC) 1 vs. PC2 and PC3 vs PC4 for the top 3,000 variable genes. The percentage of total variance assigned to each component is indicated in the axis title. **c-e**. Differential gene expression analyses contrasting BCi.NS1.1 and primary –derived HAE cultures by cell population for basal, suprabasal, intermediate, secretory I, and secretory II cells. **c**. Volcano plots summarizing the 1,052 differentially expressed genes (DEGs) calculated from pseudobulk profile contrasts, defining significance as an adjusted p-value (Benjamini-Hochberg) < 0.05, log2FC > 1, and per-cell population expression in > 10% of cells. TERT and ISGs IFIT1, ISG15, and MX1 are annotated where significantly differentially expressed, along with the number of DEGs for each cell population **d**. An “Upset” bar plot displaying the intersection of DEGs by cell population. The majority DEGs are unique to a particular cell population, but a core set of 30 genes (outlined in gold) were similarly differentially expressed across all groups. **e**. Gene set enrichment testing using CAMERA for the top 30 C2 canonical pathways gene sets by FDR, ordered by directional p-value across cell populations. Sets which were not significantly different across conditions are shown in dark grey. A positive directional p-value represents sets which are positively differentially expressed in BCi-NS1.1-vs primary-derived HAE cultures and a negative value represents negatively differentially expressed sets. Secretory II cells in particular display an elevated interferon response signature, highlighted in red: “WP Type II Interferon Signaling”, “Reactome Interferon Alpha Beta Signaling”, “WP Immune Response to Tuberculosis”, “WP Type I Interferon Induction And Signaling During SARS-CoV-2 Infection”.

### Pseudobulk differential gene expression reveals higher interferon-stimulated gene (ISG) expression in some BCi-NS1.1 HAE culture cell populations

Despite the high degree of similarity in HAE culture cell populations across culture progenitor groups, the relatively minor gene expression differences observed between BCi-NS1.1 and primary-derived HAE culture cell populations in PCA could be biologically meaningful. Therefore, for each cell population, we conducted differential gene expression analysis directly comparing BCi-NS1.1 and primary HAE cultures. In total, we detected 1,052 genes differentially expressed in at least one cell population of those with sufficient cell numbers for testing (basal, suprabasal, intermediate, secretory I, secretory II, Fig. 3c, Supplementary Data File 1). A “core” set of 30 genes were detected as differentially expressed (same direction) in all cell populations tested (Fig. 3d). The majority of differentially expressed genes (DEGs) were detected in a single cell population (Fig. 3d). As expected, *TERT* expression was higher in all BCi-NS1.1-derived HAE culture cell populations, though it failed to meet differential expression thresholds in all but secretory I and II cells due to sparse detection typical of droplet scRNA-Seq (Fig. 3c, Fig. S2a).

To place DEGs in biological context we performed gene set enrichment testing on the C2 canonical pathway gene sets from the Molecular Signatures Database^46^. Several gene sets related to interferon (IFN) signaling were found to be enriched in BCi-NS1.1 cultures, particularly in secretory II cells (Fig. 3e, Fig. S2a). Relatedly, we observed differential expression of ISGs *MX1*, *IFIT1*, and *ISG15* in secretory II cells and *ISG15* and *MX1* expression in basal cells (indicated in Fig. 3c). Thus, while gene expression is largely similar, a relatively small number of genes are differentially expressed between BCi-NS1.1 and primary HAE culture cell populations. Among these, we observed an elevated ISG signature in some BCi-NS1.1. cell populations.

### JAK-STAT pathway activity and expression of select ISGs are higher at steady state in BCi-NS1.1 relative to primary HAE cultures, but are induced to similar levels upon IAV infection

As HAE cultures are frequently employed for infection studies by our group and others^1,6,7,14,22,47^, we sought to further characterize potential differences in ISG expression and corresponding signaling between these two culture types at steady state, in response to IFN stimulation, and in response infection with respiratory pathogens. We apically infected both types of cultures with two human respiratory pathogens: influenza A virus (IAV) and *Staphylococcus aureus*. As a positive control for ISG expression, we treated cultures basolaterally with human IFN-β. Mock-infected cultures served as controls. We measured the expression of select ISGs (identified in scRNA-Seq analyses) by RT-qPCR and by western blot (Fig. 4a-f). We also measured phosphorylated STAT1 to assess JAK-STAT pathway activity. Mock-treated BCi-NS1.1-derived HAE cultures showed higher expression of ISGs examined (MX1, IFIT1, ISG15) than primary-derived HAE cultures, as observed in our scRNA-Seq analysis (Fig. 3c, e). This difference was statistically significant for MX1 at both RNA and protein levels (Fig. 4a-b). Concomitantly, phosphorylated STAT1 was elevated in BCi-NS1.1-derived cultures at steady-state (Fig. 4b).

**Fig. 4:**
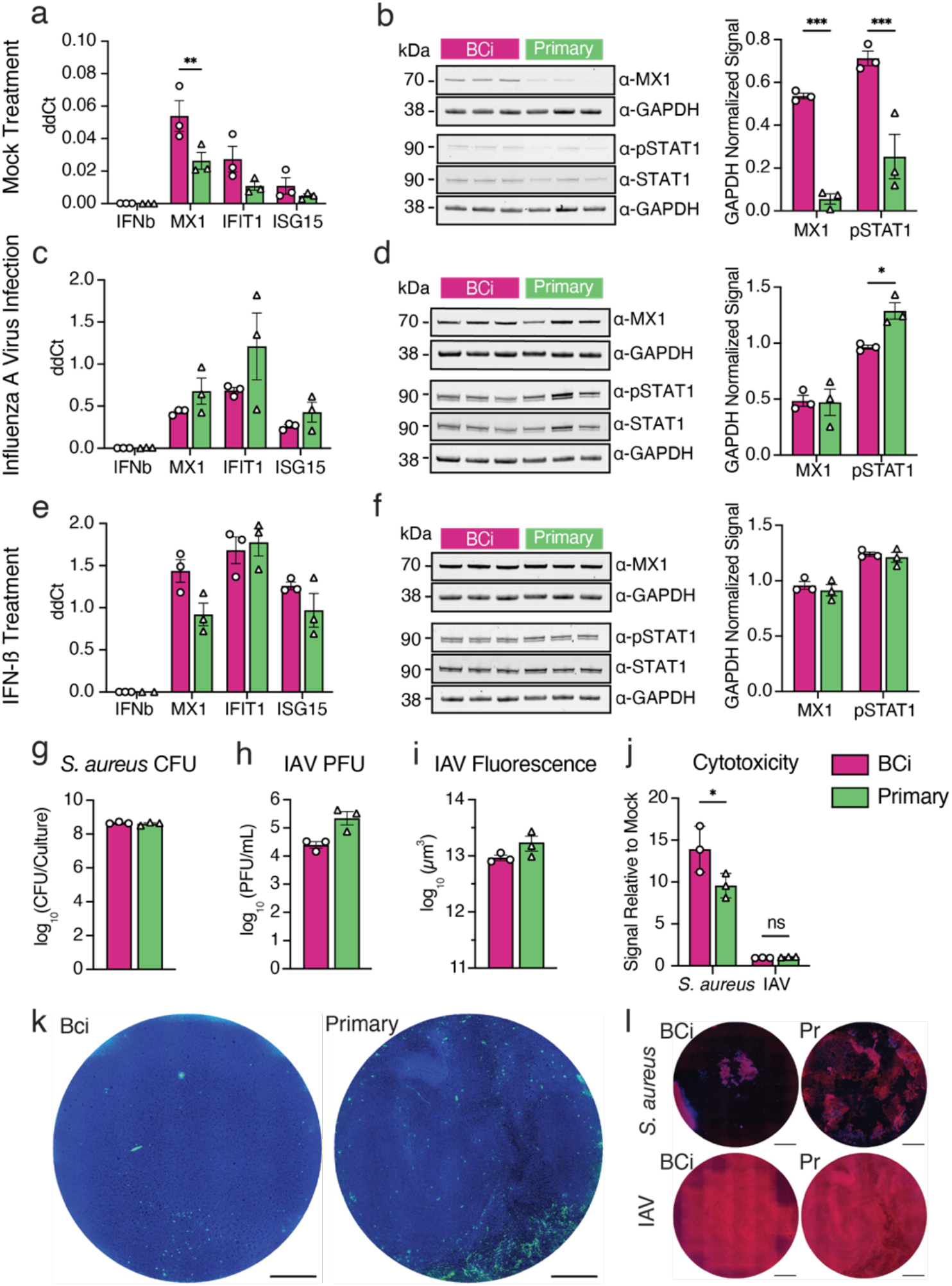
Infection of BCi-NS1.1- and primary-derived HAE cultures with IAV and *S. aureus*. **a, c, e**. RNA levels of MX1, IFN-b, IFIT1 and ISG15 were measured using RT-qPCR at steady-state (a), after IAV infection (c) and after IFN-b treatment. **b, d, f**. Representative western blots for MX1 and phosphorylated STAT1 (p-STAT1), along with GAPDH controls for mock treated (b), influenza A virus (IAV)-infected (d) and IFN-b-treated (f) cultures, and quantification of western blot band intensity from three individual blots for each condition. **g**. *S. aureus* colony-forming units (CFU) after 18h of infection **h**. IAV plaque-forming units (PFU) after 24h of infection **i**. influenza A virus-infected cultures were fixed and stained with an antibody against influenza virus-NP, then infection was quantified as total NP positive area. **j**. LDH release was quantified after influenza A virus or *S. aureus* infection and plotted as a fold change of LDH release over mock infected cultures. **k**. Top-down images of representative IAV-infected HAE cultures, showing DAPI (blue) and influenza virus-NP (green). Scale bars = 1mm. **l**. Top-down images of representative *S. aureus* and IAV-infected HAE cultures, showing Phalloidin (red) and DAPI (blue). Scale bars = 1mm.

We next sought to determine whether these differences in steady-state ISG expression result in functional differences upon infection with common respiratory pathogens. As expected, we observed mRNA upregulation of ISGs (MX1, IFIT1 and ISG15) upon influenza A virus infection (Fig. 4c, d) and IFN-ß treatment (Fig. 4e, f). Interestingly, despite the higher baseline ISG expression in BCi-NS1.1-derived cultures at steady-state, ISG RNA expression was upregulated to similar levels in in primary cell-derived HAE cultures and BCi-NS1.1-derived cultures. MX1 protein levels, while different between culture types at baseline (Fig. 4b), were also induced to similar levels upon IAV infection (Fig. 4d) and IFN-ß treatment (Fig. 4f). Finally, phosphorylated STAT1 levels, while higher in BCi-NS1.1-derived cultures at steady state (Fig. 4b), exhibited an inverse pattern upon IAV infection (Fig. 4c). No difference in phosphorylated STAT1 levels was observed upon IFN-ß stimulation (Fig. 4f). We did observe more variable ISG upregulation (RNA) in primary-derived HAE cultures than in BCi-NS1.1 cell-derived HAE cultures (Fig. 4c, e), possibly due to donor-associated variation.

### Primary and BCi-NS1.1-derived HAE cultures exhibit similar pathogen loads and tissue damage upon infection

We next determined whether the observed increased baseline ISG expression in BCi-NS1.1-derived HAE cultures affects pathogen loads upon infection. Thus, at 24 or 18 hours post-infection, respectively, we measured IAV plaque-forming units (PFU) or *S. aureus* colony-forming units (CFU) in apical washes from BCi-NS1.1- and primary cell-derived HAE cultures. In addition, we measured pathogen-elicited tissue damage by lactate dehydrogenase (LDH) release and fluorescence microscopy. We found that *S. aureus* loads were equal between both culture types (Fig. 4g). IAV loads trended higher in primary cell-derived HAE cultures (Fig. 4h, I, k, Fig. S4a), but this difference was not statistically significant. These modest differences may be due to the elevated steady-state ISG expression observed in BCi-NS1.1-derived cultures (Fig. 4a-b), but further investigation would be needed to confirm this correlation. IAV infection resulted in low-level LDH release with no detectable differences across culture types (Fig. 4j). *S. aureus* infection resulted in considerable LDH release, with higher levels in BCi-NS1.1-derived cultures. These patterns were consistent with corresponding fluorescence microscopy of infected cultures to assess cellular damage (Fig. 4l, Fig. S4b). Taken together, these results demonstrate that, despite some minor variations, BCi-NS1.1- and primary-derived HAE cultures behave similarly upon infection with two major respiratory pathogens, influenza A virus and *S. aureus*.

## Conclusions

Combining scRNA-Seq, immunohistochemistry, and functional analyses, we have shown that HAE cultures derived from the hTERT-expressing precursor cell line BCi-NS1.1 exhibit similar cell type composition and similar cell type-associated gene expression to HAE cultures derived from primary cells. While we observed evidence of heightened JAK/STAT pathway activity and corresponding increased baseline expression of several ISGs in BCi-NS1.1-derived HAE cultures at steady-state, both culture types induced ISG expression to similar levels upon infection with IAV or upon IFN-treatment. Taken together, these results further support BCi-NS1.1-derived HAE cultures as a relevant model system that recapitulates airway epithelial biology similarly to primary HAE cultures. These cultures are of particular benefit in experiments that demand large quantities of HAE cultures/replicates beyond those available from primary cell sources. Additionally, the extended passage capacity of BCi-NS1.1 cells enables sophisticated genetic engineering techniques (with corresponding antibiotic selection) for mechanistic studies of airway cell function and disease^26^ that can be technically challenging or infeasible in primary cells.

Despite overall similarities, we did observe modest differences between culture types. These differences may stem from donor-to-donor variation as all BCi-NS1.1 cultures were necessarily derived from the same donor while primary cultures were derived from three distinct donors, and/or directly from engineered expression of hTERT in BCi-NS1.1 cells. Future studies in which hTERT expression is engineered in multiple independent primary cell sources may disentangle these mechanisms. Further, determining the specific effects of hTERT in HAE cultures will be important in establishing and interpreting hTERT-engineered culture models of specific demographic groups or disease states for which primary HAE culture systems have been proven highly informative, such as cystic fibrosis,^48^ COPD,^49^ and asthma^50^.

## Acknowledgements

We thank Dr. Ronald Crystal for his kind gift of BCi-NS1.1 cells. We thank Dr. Margaret Scull for critical reading of the manuscript and helpful suggestions. We also thank all Dittmann and Rosenberg lab members for their thoughts and input on this project. Research was partially supported by the following grants from NIH/NIAID: R01AI143639 to M.D., AI137336 to V.J.T., R01 AI151029 to B.R.R. and U01 AI150748 to B.R.R. Work was further supported by The Vilcek Institute of Graduate Biomedical Sciences, and by NYU Grossman School of Medicine Startup funds. Both the Experimental Pathology Research Laboratory and the NYU Genome Technology Core are supported by NYU Cancer Center support grant P30CA016087 and by NYU Langone’s Laura and Isaac Perlmutter Cancer Center. Research reported in this paper was supported by the Office of Research Infrastructure of the National Institutes of Health under award number S10OD026880. The content is solely the responsibility of the authors and does not necessarily represent the official views of the National Institutes of Health.

## Code availability

All analysis code generated for this study is available on GitHub at https://github.com/BradRosenbergLab/airwayepithelialculturecomparison.

## Author contributions

**Rachel A. Prescott**: Conceptualization, Methodology, Validation, Formal analysis, Investigation, Writing – Original draft, Visualization **Alec Pankow**: Conceptualization, Methodology, Software, Validation, Formal analysis, Investigation, Data curation, Writing – Original draft, Visualization **Maren deVries**: Conceptualization, Methodology, Investigation, Writing – Review & editing **Keaton Crosse**: Conceptualization, Methodology, Validation, Formal analysis, Investigation, Writing – Review & editing, Visualization **Roosheel S. Patel** Methodology, Software **Mark Alu** Methodology, Investigation **Cynthia Loomis** Methodology, Supervision **Victor Torres** Resources, Writing – Review & editing, Supervision, Funding acquisition **Sergei Koralov** Resources, Writing – Review & editing, Supervision, Funding acquisition **Ellie Ivanova** Methodology, Investigation, Writing – Review & editing **Meike Dittmann** Conceptualization, Methodology, Resources, Writing – Original draft, Supervision, Funding acquisition **Brad R. Rosenberg** Conceptualization, Methodology, Resources, Writing – Original draft, Supervision, Funding acquisition

## Competing interests

The authors declare no competing interests.

## Notes

### Competing Interest Statement

The authors have declared no competing interest.

## References

1. Davis, A. S. et al. Validation of Normal Human Bronchial Epithelial Cells as a Model for Influenza A Infections in Human Distal Trachea. Journal of Histochemistry and Cytochemistry 63, 312–328 (2015).

2. Gray, T. E., Guzman, K., William Davis, C., Abdullah, L. H. & Nettesheim, P. Mucociliary Differentiation of Serially Passaged Normal Human Tracheobronchial Epithelial Cells. Am. J. Respir. Cell Mol. Biol. 14, 104–112 (1996).

3. Garcıá, S. R. et al. Novel dynamics of human mucociliary differentiation revealed by single-cell RNA sequencing of nasal epithelial cultures. Development (Cambridge) 146, (2019).

4. Wang, G. et al. Characterization of an immortalized human small airway basal stem/progenitor cell line with airway region-specific differentiation capacity. Respir Res 20, 1–14 (2019).

5. Walters, M. S. et al. Generation of a human airway epithelium derived basal cell line with multipotent differentiation capacity. Respir Res 14, 26–30 (2013).

6. Vaidyanathan, S. et al. High-Efficiency, Selection-free Gene Repair in Airway Stem Cells from Cystic Fibrosis Patients Rescues CFTR Function in Differentiated Epithelia. Cell Stem Cell 26, 161–171.e4 (2020).

7. Kelly, J. N. et al. Comprehensive single cell analysis of pandemic influenza A virus infection in the human airways uncovers cell-type specific host transcriptional signatures relevant for disease progression and pathogenesis. Front Immunol 13, (2022).

8. Knight, D. A. & Holgate, S. T. The airway epithelium: Structural and functional properties in health and disease. Respirology 8, 432–446 (2003).

9. Eon Kuek, L. & Lee, R. J. First contact: the role of respiratory cilia in host-pathogen interactions in the airways. Am J Physiol Lung Cell Mol Physiol 319, 603–619 (2020).

10. Branchfield, K. et al. Pulmonary neuroendocrine cells function as airway sensors to control lung immune response. Science (1979) 351, 707–710 (2016).

11. Hewitt, R. J. & Lloyd, C. M. Regulation of immune responses by the airway epithelial cell landscape. Nat Rev Immunol 21, 347–362 (2021).

12. Plasschaert, L. W. et al. A single-cell atlas of the airway epithelium reveals the CFTR-rich pulmonary ionocyte. Nature 560, 377–381 (2018).

13. Ualiyeva, S. et al. Airway brush cells generate cysteinyl leukotrienes through the ATP sensor P2Y2. Sci. Immunol 5, (2020).

14. Montoro, D. T. et al. A revised airway epithelial hierarchy includes CFTR-expressing ionocytes. Nature 560, 319–324 (2018).

15. Ibricevic, A. et al. Influenza Virus Receptor Specificity and Cell Tropism in Mouse and Human Airway Epithelial Cells. J Virol 80, 7469–7480 (2006).

16. Wu, C. T. et al. SARS-CoV-2 replication in airway epithelia requires motile cilia and microvillar reprogramming. Cell 186, 112–130.e20 (2023).

17. Gagneux, P. et al. Human-specific Regulation of α2-6-linked Sialic Acids. Journal of Biological Chemistry 278, 48245–48250 (2003).

18. Shinya, K. et al. Influenza virus receptors in the human airway. (2006).

19. Dohrman, A. et al. Mucin gene MUC 2 and MUC 5AC upregulation by Gram-positive and Gram-negative bacteria. Biochim Biophys Acta 1406, (1998).

20. Chen, R. et al. Nontypeable Haemophilus influenzae lipoprotein P6 induces MUC5AC mucin transcription via TLR2-TAK1-dependent p38 MAPK-AP1 and IKKβ-IκBα-NF-κB signaling pathways. Biochem Biophys Res Commun 324, 1087–1094 (2004).

21. Kim, Y. O., Jung, M. J., Choi, J. K., Ahn, D. W. & Song, K. S. Peptidoglycan from staphylococcus aureus increases MUC5AC gene expression via RSK1-CREB pathway in human airway epithelial cells. Mol Cells 32, 359–365 (2011).

22. Ravindra, N. G. et al. Single-cell longitudinal analysis of SARS-CoV-2 infection in human airway epithelium identifies target cells, alterations in gene expression, and cell state changes. PLoS Biol 19, (2021).

23. Chu, H. W. et al. CRISPR-Cas9-mediated gene knockout in primary human airway epithelial cells reveals a proinflammatory role for MUC18. Gene Ther 22, 822–829 (2015).

24. Zhou, H. et al. POU2AF1 Functions in the Human Airway Epithelium To Regulate Expression of Host Defense Genes. The Journal of Immunology 196, 3159–3167 (2016).

25. Iverson, E. et al. Membrane-Tethered Mucin 1 Is Stimulated by Interferon and Virus Infection in Multiple Cell Types and Inhibits Influenza A Virus Infection in Human Airway Epithelium. mBio 13, (2022).

26. Song, D. et al. MUC5B mobilizes and MUC5AC spatially aligns mucociliary transport on human airway epithelium. (2022).

27. Vries, M. de et al. A Comparative Analysis of SARS-CoV-2 Antivirals Characterizes 3CL pro Inhibitor PF-00835231 as a Potential New Treatment for COVID-19. (2021).doi:10.1128/JVI

28. Mimitou, E. P. et al. Multiplexed detection of proteins, transcriptomes, clonotypes and CRISPR perturbations in single cells. Nat Methods 16, 409–412 (2019).

29. Stoeckius, M. et al. Cell Hashing with barcoded antibodies enables multiplexing and doublet detection for single cell genomics. Genome Biol 19, (2018).

30. Hao, Y. et al. Integrated analysis of multimodal single-cell data. Cell 184, 3573–3587.e29 (2021).

31. McGinnis, C. S., Murrow, L. M. & Gartner, Z. J. DoubletFinder: Doublet Detection in Single-Cell RNA Sequencing Data Using Artificial Nearest Neighbors. Cell Syst 8, 329–337.e4 (2019).

32. Hafemeister, C. & Satija, R. Normalization and variance stabilization of single-cell RNA-seq data using regularized negative binomial regression. Genome Biol 20, (2019).

33. Tirosh, I. et al. Dissecting the multicellular ecosystem of metastatic melanoma by single-cell RNA-seq.

34. Rotta, R. & Noack, A. Multilevel local search algorithms for modularity clustering. In ACM Journal of Experimental Algorithmics 16, (Association for Computing Machinery, 2011).

35. Zappia, L. & Oshlack, A. Clustering trees: a visualization for evaluating clusterings at multiple resolutions. Gigascience 7, (2018).

36. Carraro, G. et al. Transcriptional analysis of cystic fibrosis airways at single-cell resolution reveals altered epithelial cell states and composition. Nat Med 27, 806–814 (2021).

37. Robinson, M. D., McCarthy, D. J. & Smyth, G. K. edgeR: A Bioconductor package for differential expression analysis of digital gene expression data. Bioinformatics 26, 139–140 (2009).

38. Lun, A. T. L., McCarthy, D. J. & Marioni, J. C. A step-by-step workflow for low-level analysis of single-cell RNA-seq data [version 1; referees: 5 approved with reservations]. F1000Res 5, (2016).

39. Squair, J. W. et al. Confronting false discoveries in single-cell differential expression. Nat Commun 12, (2021).

40. Wu, D. & Smyth, G. K. Camera: A competitive gene set test accounting for inter-gene correlation. Nucleic Acids Res 40, (2012).

41. Liberzon, A. et al. Molecular signatures database (MSigDB) 3.0. Bioinformatics 27, 1739–1740 (2011).

42. Street, K. et al. Slingshot: Cell lineage and pseudotime inference for single-cell transcriptomics. BMC Genomics 19, (2018).

43. Lun, A. T. L., Richard, A. C. & Marioni, J. C. Testing for differential abundance in mass cytometry data. Nat Methods 14, 707–709 (2017).

44. Boles, B. R., Thoende, M., Roth, A. J. & Horswill, A. R. Identification of genes involved in polysaccharide-independent Staphylococcus aureus biofilm formation. PLoS One 5, (2010).

45. Başak, K., Kumbul Doguç, D., Aylak, F., Karadayı, N. & Gültekin Lütfi Kırdar Kartal, F. Effects of Maternally Exposed Food Coloring Additives on Laryngeal Histology in Rats. Journal of Environmental Pathology 33, (2014).

46. Liberzon, A. et al. The Molecular Signatures Database Hallmark Gene Set Collection. Cell Syst 1, 417–425 (2015).

47. V’kovski, P. et al. Disparate temperature-dependent virus–host dynamics for SARS-CoV-2 and SARS-CoV in the human respiratory epithelium. PLoS Biol 19, (2021).

48. Moreau-Marquis, S. et al. The F508-CFTR mutation results in increased biofilm formation by Pseudomonas aeruginosa by increasing iron availability. Am J Physiol Lung Cell Mol Physiol 295, 25–37 (2008).

49. Horndahl, J. et al. HDAC6 inhibitor ACY-1083 shows lung epithelial protective features in COPD. PLoS One 17, (2022).

50. Schindler, V. E. M. et al. Side-Directed Release of Differential Extracellular Vesicle-associated microRNA Profiles from Bronchial Epithelial Cells of Healthy and Asthmatic Subjects. Biomedicines 10, 622 (2022).

