## Supplemental Figures for "A comparative study of *in vitro* air-liquid interface culture models of the human airway epithelium evaluating cellular heterogeneity and gene expression at single cell resolution"

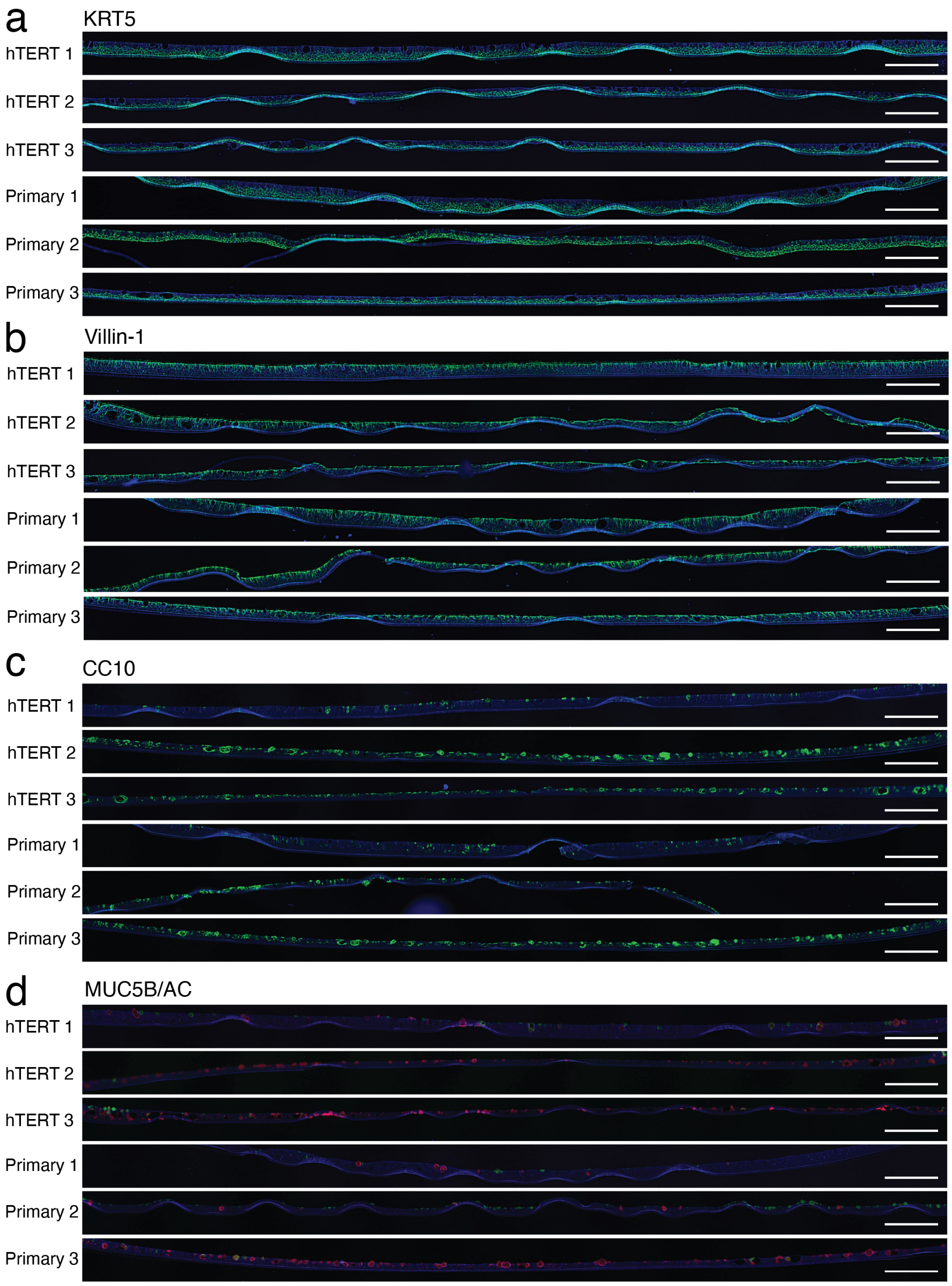


**Supplementary Figure 1: Fixed and cross-sectioned HAE culture sections at steady-state from each of the six cultures imaged for cell-type staining.** Scale bars = 200μm. Phalloidin (blue) a. KRT5 (green) b. Villin-1 (green) c. CC10 (green) d. MUC5AC (red) and MUC5B (green).

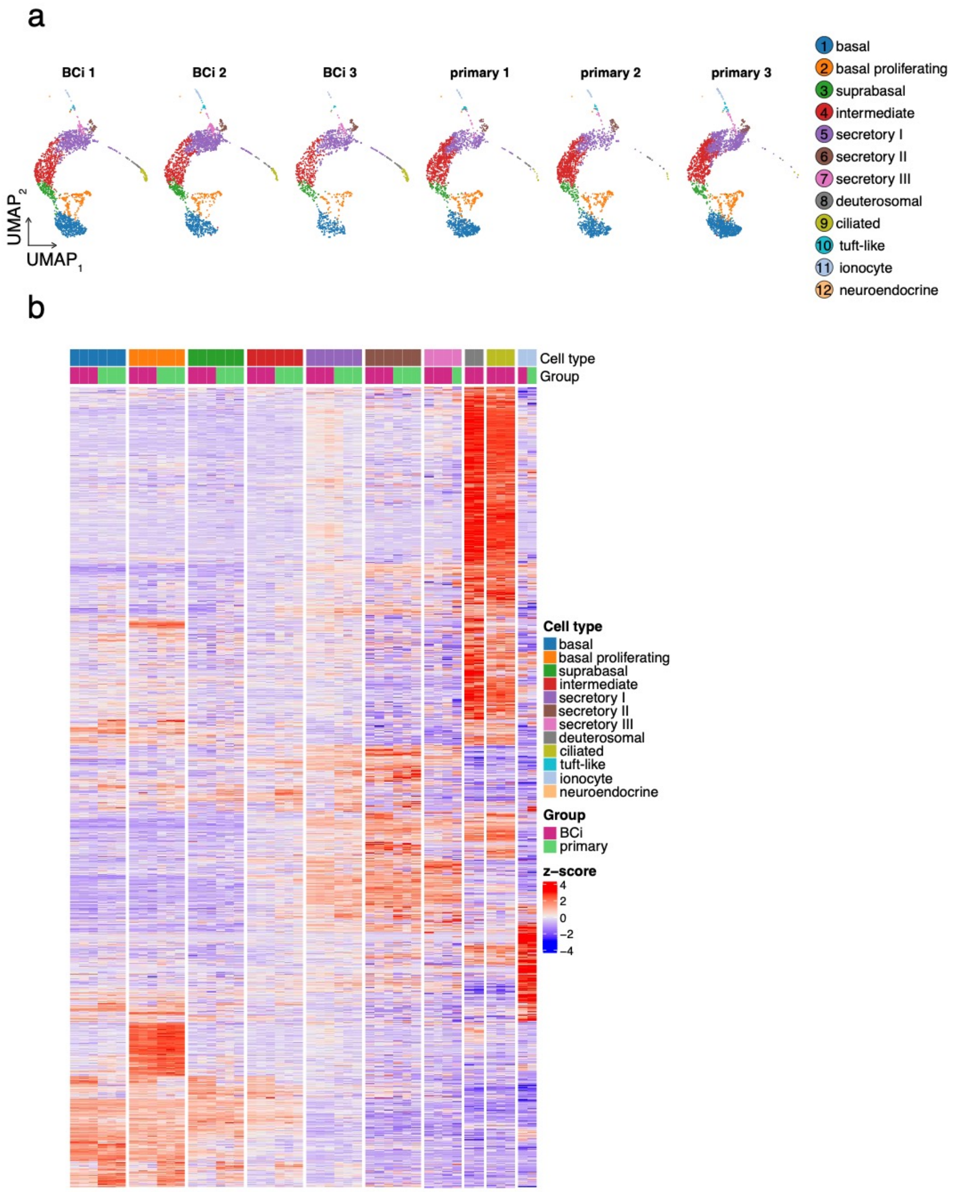


**Supplementary Figure 2: scRNA-Seq cell population assignment by replicate and marker gene expression patterns.** UMAP dimensionality reduction plots of scRNA-Seq independently normalized with scTransform (v.0.3.3) and integrated using Seurat 4.0, displaying uniformity in cell type annotation across replicates. **b.** A large heatmap of 4,001 marker genes by cell population from scRNA-seq, with rows clustered by enrichment pattern and column ordering by cell and progenitor type. For a list of all marker genes, see Supplementary Data File 2. Differentially expressed markers were identified by contrasting profiles within a cell population with the average expression of remaining cell annotations, using significance cutoffs of $Log_2FC > 2$ (positively expressed only) and an FDR of < 0.05. Clear modules of cell-type specific gene expression common in BCi-NS1.1- and primary-derived HAE cultures are apparent, particularly for proliferating, deuterosomal, ciliated cells and ionocytes.

a

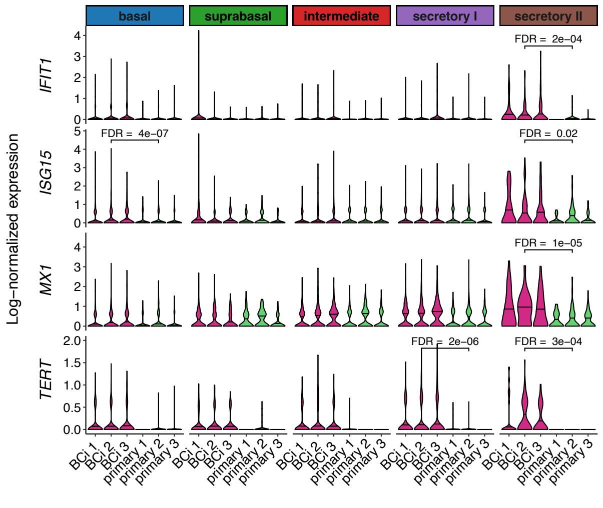


b

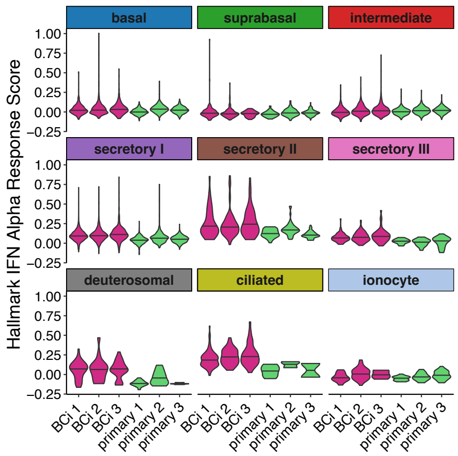


**Supplementary Figure 3: a.** Violin plots of log-normalized expression values for ISGs *IFIT1*, *ISG15*, and *MX1* along with *TERT* by cell population. Plots are annotated with the FDR from pseudobulk differential expression testing across BCi-NS1.1-derived and primary HAE where significant. **b.** Violin plots of “Hallmark Interferon Alpha Response” gene set scores at the single cell level by cell population. Low sampling of secretory III, deuterosomal, ciliated cells and ionocytes required their exclusion from pseudobulk contrasts, but they are included here for completeness.

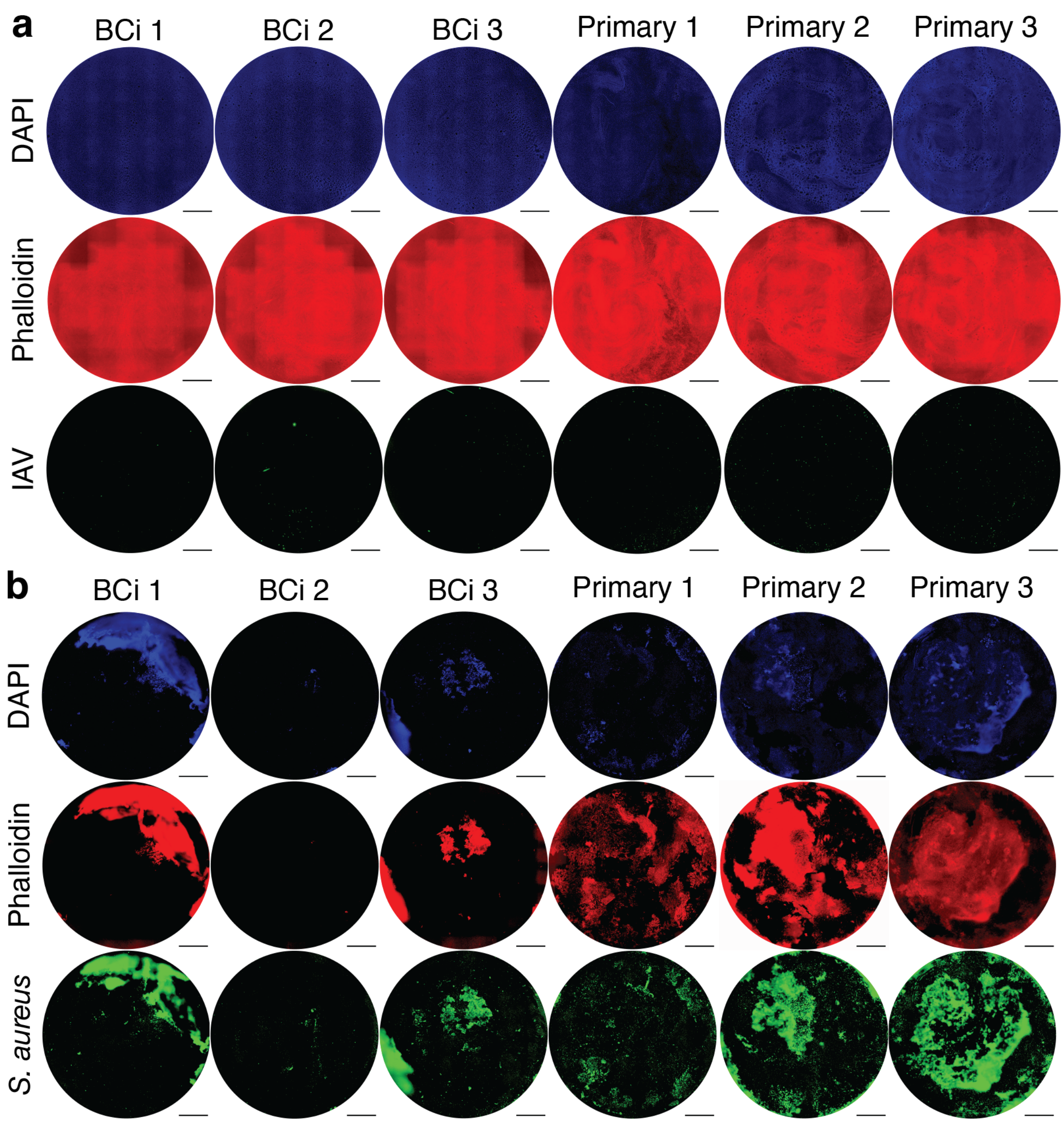


**Supplementary Figure 4:** Top-down images of HAE cultures with individual channels stained for phalloidin (red) and DAPI (blue) and infected with **a.** IAV (green) and **b.** *S. aureus* (green). Scale bars = 1mm.
